# Capturing Cardiomyocyte Cell-to-Cell Heterogeneity via Shotgun Single Cell Top-Down Proteomics

**DOI:** 10.64898/2026.03.26.714071

**Authors:** Fabio P. Gomes, Blandine Chazarin, Aleksandra Binek, Aleix Navarro Garrido, Kenneth R. Durbin, Ricard Garcia-Carbonell, Kanchan Pathak, Delaynie Brinkman, Reynaldo Magalhaes Melo, Anja Karlstaedt, Enrique Saez, Jennifer E. Van Eyk, John R. Yates

## Abstract

Individual cells exhibit distinct molecular landscapes shaped by proteins and the diverse functional repertoire of their corresponding proteoforms. These structurally diverse variants (e.g., post-translationally modified including truncated proteolyzed forms) collectively orchestrate cellular functions. However, resolving proteoform heterogeneity at single-cell (SC) resolution remains a significant analytical challenge. Here, we present a shotgun SC top-down proteomics (SC-TDP) strategy that enables direct, unbiased proteoform profiling from single cardiomyocytes. Across 13 individual cardiomyocytes isolated from mouse heart, we identified a total of 57 proteins represented by 165 distinct proteoforms, including phosphorylated, succinylated, trimethylated, truncated, amongst others. Notably, proteoform composition varied substantially among cells, revealing a previously unrecognized level of molecular heterogeneity among cardiomyocytes. Together, these findings establish SC-TDP as a powerful tool for uncovering the proteoform diversity at the SC level. Our strategy paves the way for defining functional heterogeneity in cardiac tissue with unprecedented molecular resolution, enabling direct examination of the proteoform landscape that underlies cellular identity and physiology.

## Main Text

The discovery that individual cells have unique molecular compositions, and thus, biological function, has opened new avenues for medical diagnoses and disease treatments.^1^ It is, therefore, of great interest to define “cell-to-cell” heterogeneity. Single-cell (SC) RNA sequencing reveals the complexity and variability of mRNA transcripts.^2^ However, this technology is unable to precisely define the functionality of the cells because proteins, including their modified forms (proteoforms), are the key functional players within cells. Mass spectrometry (MS)-based proteomics can comprehensively profile cellular proteins and their proteoforms with high precision. It has emerged as a powerful tool for studying these biomolecules in SCs.^1,3-5^ Bottom-up proteomics, in particular, has provided new insights into disease mechanisms and intercellular communication by identifying specific proteins involved in these processes.^6-10^ Nevertheless, this approach is limited in its ability to elucidate proteoforms representing the diverse modified protein products derived from a single gene. Because protein digestion is an integral part in bottom-up proteomics, information regarding the co-occurrence of distinct posttranslational modifications (PTMs) or sequence variations on the same protein is often lost. In addition, the protein inference problem further limits accurate proteoform characterization.^10-12^ Proteoforms can independently modulate numerous biological processes, and they may also serve as effective indicators of health and disease.^13,14^ Because top-down proteomics (TDP) is performed on intact proteins rather than digested peptides, protein structural deviations are retained.^10,11^ The characterization of proteoforms in individual cells is notoriously challenging due to extremely low input amounts, limited throughput, and the technical difficulties associated with the proteoform separation and effective top-down fragmentation.^5,10,15,16^

Despite these challenges, the field of SC-TDP has progressed steadily, albeit with relatively few publications, evolving from early proof-of-concept studies on erythrocytes^17,18^ to the development of modern high-throughput imaging^3,5^ and microfluidic platforms.^10,14^ This progression is exemplified by recent studies demonstrating the feasibility of top-down analysis in individual cells. For instance, the introduction of SC proteoform imaging MS (scPiMS) enabled intact proteoform profiling of individual cells from the rat hippocampus.^3^ Another example includes a top-down analysis of single HeLa cells employed on-capillary cell lysis and identified ∼50 proteoforms using a manual sample-loading approach.^14^ Additionally, TDP was applied to investigate heterogeneity of large proteoforms in single muscle fibers. Notably, using a Bradford protein assay, the authors estimated the total protein content of individual fibers (∼100 ng), with lengths ranging from ∼1500 to 2500 µm.^10^

Here, we expand on this SC-TDP evolution by introducing a high-resolution proteoform analysis approach using liquid chromatography-mass spectrometry (LC–MS). Specifically, we demonstrate that using electron-transfer higher-energy collision dissociation (EThcD) and higher-energy collision dissociation (HCD) within the same scan effectively improves both proteoform coverage and protein backbone fragmentation. This method provides a simple, robust, fast, ultrasensitive, and high-throughput platform for efficiently analyzing proteoforms in individual cardiomyocytes isolated from mouse hearts.

The sample preparation and data acquisition/analysis are described in detail in the *Supporting Information*. Briefly, individual cardiomyocytes were collected onto a 384-well plate using a CellenONE X1 device after isolation from a mouse heart tissue, cells were directly lysed onto the plate and placed into the LC-MS system for analysis, eliminating sample transfer/handling. As sample amount and proteoform content in SCs are extremely limited, minimizing losses during sample handling and preparation is critically important. This leaves essentially no opportunity for additional preparation or purification steps. We optimized the lysis buffer to simultaneously disrupt the cell, extract proteins, and denature them in a single step. To meet these requirements, our lysis buffer was primarily composed of organic solvents, including trifluoroethanol (TFE) and dimethyl sulfoxide (DMSO) which should preserve proteoform integrity and promote efficient TDP fragmentation of the protein backbone.

Using a mixture of 6 standard proteins (∼9–68 kDa), we optimized our LC-MS workflow and verified that the lysis buffer did not cause protein precipitation during analysis. Although the protein standards were diluted in a lysis buffer rich in organic solvents, we were able to detect 5 of the 6 proteins **(Support Information Figure S1)**. The largest protein (∼68kDa) was not detected, likely reflecting the limitations of the mass spectrometer in analyzing intact proteins of this size.

Next, we validated our SC-TDP workflow using a diluted bulk cardiomyocyte sample as a technical control. Homogenized and solubilized cardiomyocyte (isolated from mouse left ventricle heart tissue) was analyzed in triplicate (10 ng total protein). The chromatographic profile of the bulk cardiomyocyte sample is shown in **Figure S2**. The myosin regulatory light chain 2 (ventricular/cardiac muscle isoform, MLC-2) proteoforms bearing trimethylation at A1 (magenta) and ATP synthase F (0) complex subunit E, mitochondrial (ATP5ME) in blue were consistently detected across all three replicates **(Figure S2)**. Retention times were highly reproducible across runs, supporting robust chromatographic performance. In total, we identified 41 proteins and 83 proteoforms **(Table S1)**. The mass distribution of the identified proteoforms ranged from ∼2 to 22 kDa (bulk tissue, **Figure S3**). As illustrated in **Figure S4**, overlap across technical replicates supports reproducibility at multiple levels. Protein identifications exhibit greater overlap than proteoform identifications, consistent with the challenges of top-down data acquisition. The observed overlaps indicates reproducible detection of a core set of species in discovery TDP,^19^ with the remaining discrepancy attributable to stochastic sampling and undersampling effects inherent to MS/MS fragmentation.

Finally, we specifically analyzed 13 individual cardiomyocytes that ranged between 61.04 and 67.92 µm in length. Typically, mouse cardiomyocytes range from 35-147 µm^20^, but we limited our cells to be approximately the same size to reduce any impact due to differences in starting proteoform quantities. Our SC-TDP workflow is shown in **Figure S5**. A key feature of our approach is the use of TFE and DMSO in the lysis buffer. While TFE efficiently solubilizes proteins^21^, DMSO enhances solubility^22^, induces denaturation^23^, and improves MS sensitivity.^24^ This combination is particularly well-suited for SC-TDP applications with extremely limited protein amounts and sample sizes. Another important feature of our SC-TDP workflow is the use of a chromatographic column with a small particle size (2.7 µm) that enhanced separation efficiency.^15^ Spectral quality benefited from a high number of µscans at the MS level while increasing the RF levels further improved the transmission of high m/z ions. Our EThcD strategy evolved from our previous publications (Gomes *et al*.).^25-27^ The data was acquired using EThcD and HCD in a single acquisition method. EThcD outperformed HCD **(Figure S6)**. However, the use of HCD was necessary to mitigate the long duty cycle, as HCD scans are performed faster than EThcD scans. In numerous cases, we achieved sufficient fragmentation along the protein backbone, enabling confident identifications and precise localization of PTMs. Analysis of 13 individual cells yielded 57 proteins and 165 proteoforms **(Table S2)**, with the identified biomolecules largely overlapping those observed in the bulk sample **(Figures S7). Figure 1** demonstrates the reproducibility of chromatographic profiles across 13 cells, confirming the robustness of the SC-TDP workflow. Notably, our method enabled detection of a consistent subset of MLC-2 proteoform (magenta) bearing A1 trimethylation and ATP5ME (blue) across all 13 individual cardiomyocytes, in agreement with the bulk sample. **Figure S8** illustrates the mass distribution of identified proteoforms (∼2 - 22 kDa), consistent with the bulk model. Interestingly, the highest number of proteins (36) and proteoforms (70) were observed in the cardiomyocyte measuring 67.92 µm while the lowest numbers (14 proteins and 18 proteoforms) were detected in the cell 64.26 µm **(Table S2). Figures 2A-B** illustrates the overlap of proteins and proteoforms across cardiomyocytes, respectively. At the protein level **(Figure 2A)**, cardiomyocytes exhibit extensive overlap, reflecting a highly conserved core proteome. In contrast, the high proportion of unique proteoforms in each cell **(Figure 2B)** reveals substantial heterogeneity among individual cardiomyocytes. These findings indicate that proteoform measurements capture subtle cell-to-cell variation masked at the protein level and demonstrate the capacity of SC-TDP to resolve functional diversity. Such heterogeneity suggests that individual cardiomyocytes exist in distinct molecular configurations, which contribute to specialized roles in cardiac physiology and disease. Several identified proteoforms have direct cardiac relevance. For example, sarcomeric proteins, such as MLC-2 and mitochondrial proteins, including cytochrome c oxidase subunit 6C (COX6c) and cytochrome bc-1 complex subunit 7 (Qcr7), were characterized. MLC-2, a key contractile protein essential for heart development and function^28^, exhibited multiple proteoforms. For instance, MLC-2 was trimethylated at N-terminus A1 **(Figure 3A)**, with precise high-resolution mass measurements confirming the 42 Da shift as trimethylation rather than acetylation. Evidence for this modification was further supported by the loss of the *b*-ion series (*b*14-*b*23) when the isobaric acetyl moiety (42.0106 Da) was computationally added to the N-terminus or trimethyl group is removed from A1 residue **(Figures 3B-C)**. Here, we provide the first evidence of MLC-2 trimethylation in single cardiomyocytes from mouse. The functional consequences of this modification, however, remain to be defined. MLC-2 was also found to harbor both N-terminal A1 trimethylation and T51 phosphorylation within the same protein backbone, each unambiguously localized by EThcD fragmentation **(Figure 3D)**. Notably, T51 phosphorylation generated diagnostic ions (e.g., *b*53 and *c59*) **(Figure 3D)**, providing definitive site-specific evidence of this modification. The diagnostic ions (*b*53 and *c59*) were further validated using TDValidator **(Figure 3D)**. Additional evidence for T51 phosphorylation is that removing the phosphoryl group from the MCL-2 sequence leads to the disappearance of the diagnostic ions and appearance of the *b*54 ion **(Figure S9)**. These observations highlight the capability of SC-TDP to resolve complex combinatorial modifications on a critical cardiac protein, providing insights into its functional diversity at the SC level. Importantly, phosphorylation of MLC-2 is critical for regulating myosin-actin interactions. It modulates contractile activity and thereby controls muscle contraction and cytoskeletal dynamics.^29,30^ In addition, N-terminal truncation at residue I10 produced a proteoform lacking the first 10 amino acids **(Figure 3E)**. The consequences of N-terminal truncation of MLC-2 remain poorly understood, but in general, proteolytic cleavage is able to generate distinct proteoforms playing critical regulatory roles in signal transduction and cellular function.^31^ Myosin light chain 3 (MYL3) is essential for muscle contraction, forming a complex with the myosin heavy chain to facilitate actin filament sliding.^32^ In our analysis, MYL3 was detected in a trimethylated form **(Figure S10A)**. A1 trimethylation was confidently assigned in the MYL3 proteoform based on the presence of diagnostic ions (*b6-b30, c12*, and *c78*). Analogous to MCL-2, A1 trimethylation is required for the generation of these diagnostic ions in MYL-3 **(Figure S10B)**. Mitochondrial dysfunction contributes to cardiomyocyte death and heart failure.^33,34^ Proper mitochondrial function is essential for sustaining cellular energy supply, particularly in tissues with high energetic demand such as the heart.^35^ COX6c, a critical monomeric subunit of the terminal mitochondrial respiratory chain enzyme, has been associated with hypercholesterolemia and atherosclerosis.^36^ Extensive EThcD fragmentation of the COX6c backbone **(Figure S11)** enabled unambiguous localization of its N-terminal acetylation. Although N-terminal acetylation is a widespread modification among eukaryotic proteins, its functional significance within mitochondria remains poorly understood.^37^ To our knowledge, this study provides the first evidence of N-terminal acetylation on COX6c; however, the functional consequences of this modification remain to be elucidated. The cytochrome bc1 complex (ubiquinol–cytochrome *c* oxidoreductase) is a key component of the mitochondrial inner membrane respiratory chain that catalyzes electron transfer from ubiquinol to cytochrome *c*.^38^ This complex is also known to generate superoxide, a reactive species that functions as an important signaling molecule at physiological concentrations.^39^ Gene inactivation studies have shown that a functional bc1 complex cannot be assembled in the absence of the Qcr7 subunit. Moreover, the N-terminal region of this subunit appears to be critical for proper assembly of the cytochrome bc1 complex.^38^ We identified a Qcr7 proteoform with K11 succinylation **(Figure S12)**. This PTM can markedly affect protein function. Its role in the heart, particularly in the context of heart failure and myofibrillar mechanics, is poorly understood.^40^

**Figure 1.**
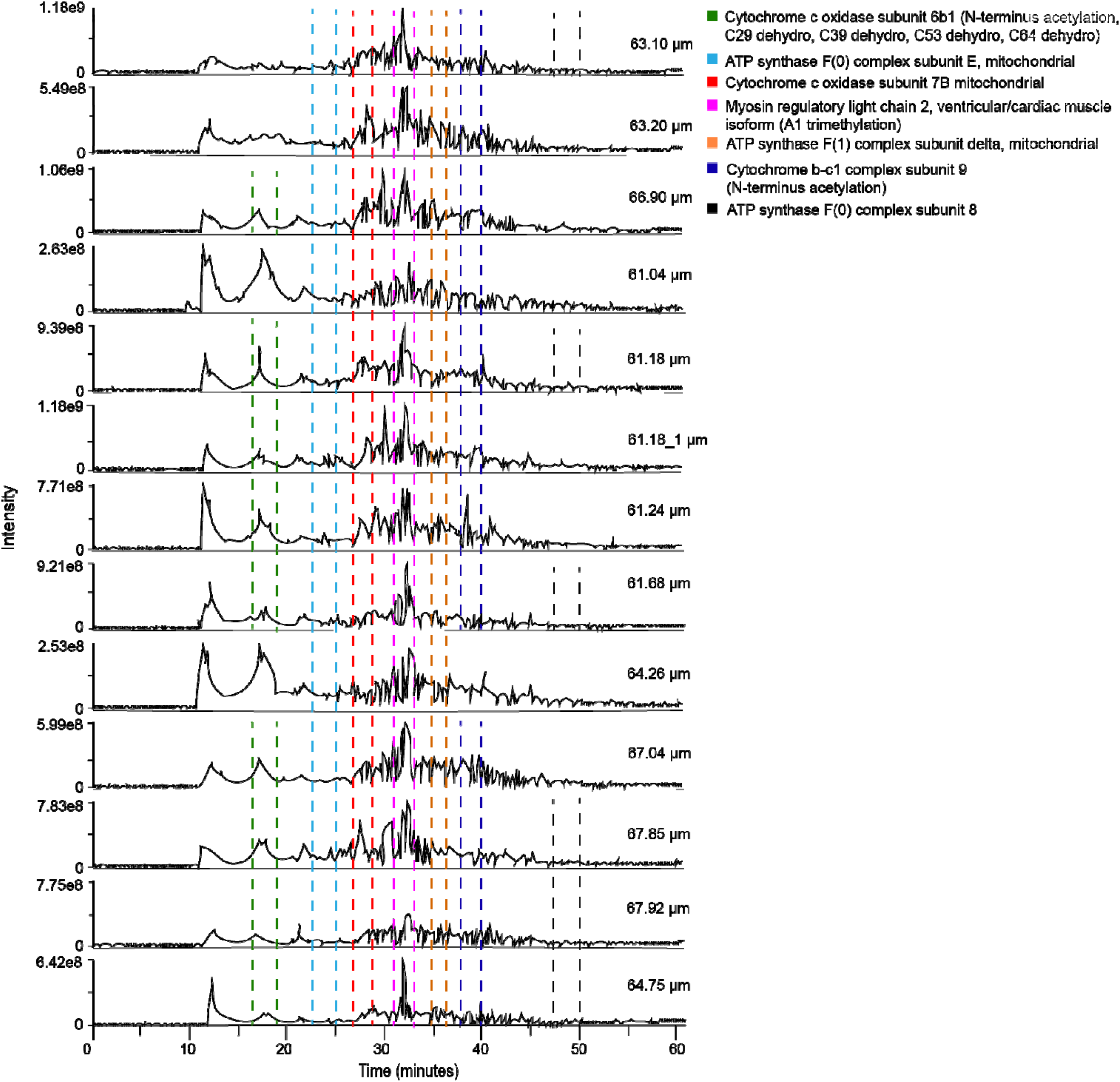
Total ion chromatograms (TICs) of 13 individual cardiomyocytes ranging from 61.04 to 67.92 µm in length. The time ranges in which each of the two selected proteins or proteoforms were detected are highlighted using different colors.

**Figure 2.**
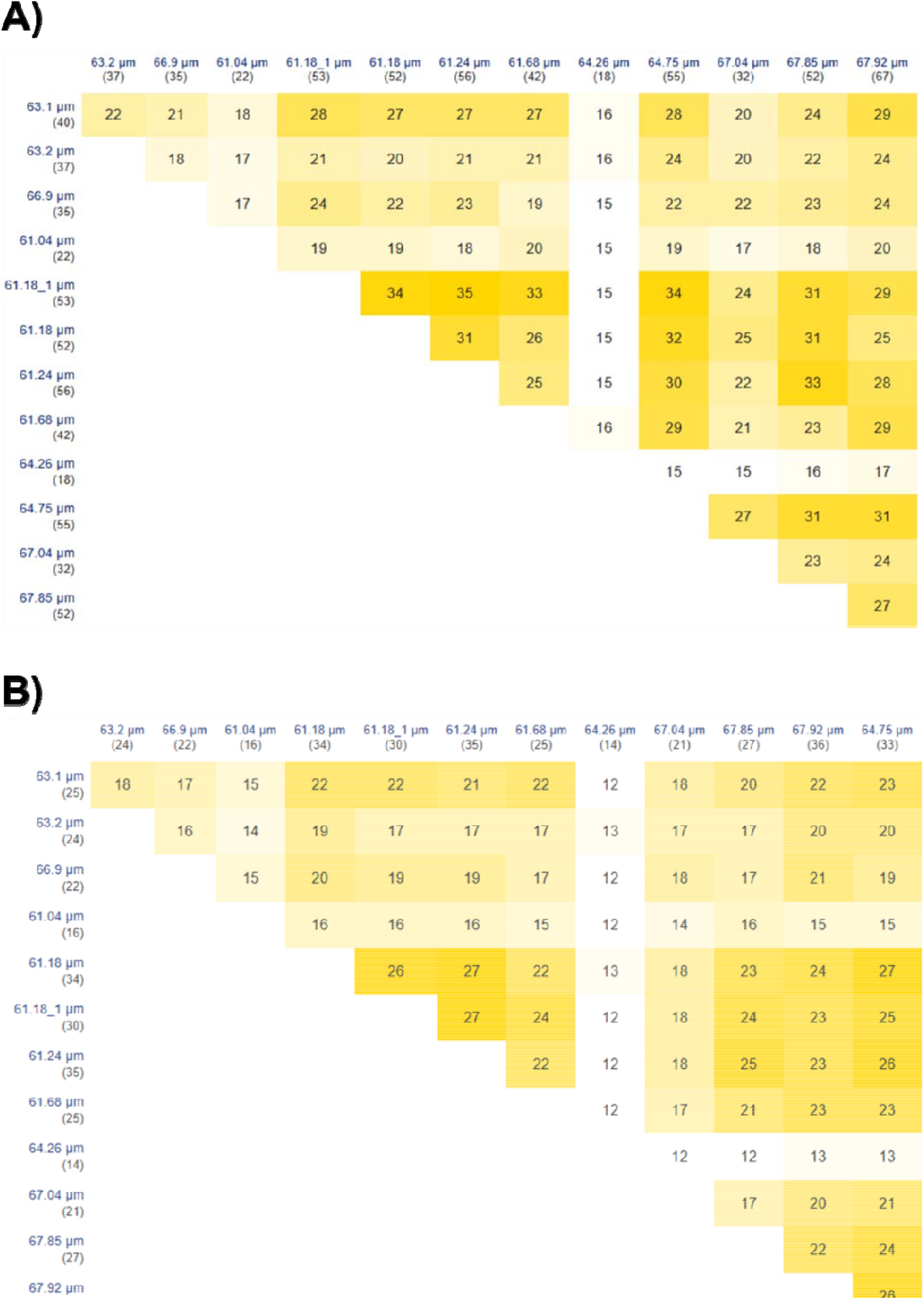
Overlap of proteins and proteoforms across 13 individual cells, each identified by its cell size (μm), with the total number of proteins or proteoforms detected shown in parentheses. The heatmaps use a light-to-dark yellow scale to indicate the degree of overlap in protein or proteoform identifications among individual cardiomyocytes. While lighter shades represent lower overlap, darker shades indicate higher overlap. **A)** Overlap of proteins across the 13 individual cardiomyocytes. Proteins were matched based on accession numbers. **B)** Overlap of proteoforms across the 13 individual cardiomyocytes. Proteoforms were compared based solely on protein descriptions and associated modifications, so the identification numbers may not represent the total number of proteoforms listed in **Table S2**.

**Figure 3.**
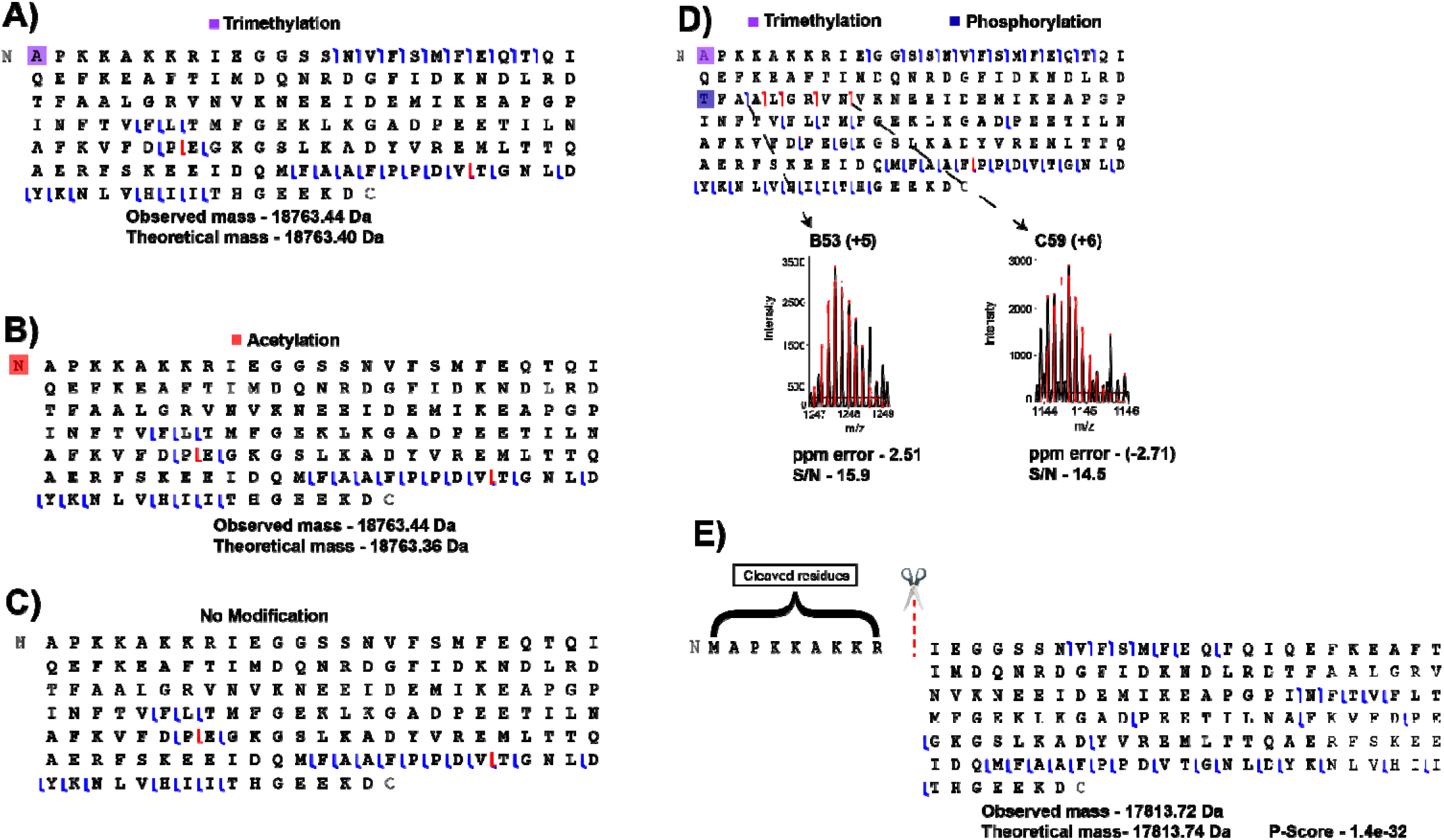
Fragmentation maps of novel MLC-2 proteoforms. **A)** MLC-2 proteoform with trimethylation at residue A1. **B)** MLC-2 proteoform with N-terminus acetylation. **C)** MLC-2 proteoform without PTMs. **D)** MLC-2 proteoform carrying trimethylation at A1 and phosphorylation at residueT51. **E)** Truncated MLC-2 proteoform. All are novel and not previously reported with exception of phosphorylation T51.

In summary, our shotgun SC-TDP strategy represents a major advance, enabling the identification and characterization of novel proteoform signatures in single cardiomyocytes. We unambiguously characterized numerous cardiac-relevant modifications, including acetylation, phosphorylation, trimethylation, and truncations. We provide evidence of succinylation in cardiomyocytes. We confidently assigned multiple PTMs in a single protein. The functional importance of combinatorial PTMs has been well documented across numerous biological processes.^41^ Expanding the delta search to 200 Da revealed many previously unknown structural deviations that may have biological significance. This threshold was selected to account for the potential incorporation of up to two phosphorylation events.^42^ This work provides the first insights into the proteoform landscape and heterogeneity of single cardiomyocytes. It offers a powerful resource for understanding cardiovascular biology at the SC level. Notably, we report the first identification of MLC-2 concurrently harboring trimethylation and phosphorylation on a single protein backbone. We also report MYL3 trimethylation. Moreover, we provide the first evidence of Qcr7 succinylation in single cardiomyocytes. While the functional consequences of most modifications remain unclear, the resulting proteoforms, may serve as potential biomarkers for cardiac disease. Myosin light chain proteoforms capture dynamic sarcomeric remodeling and contractile regulation, providing mechanistic insight into cardiac dysfunction.^43,44^ This molecular resolution enables interrogation of disease processes and regulatory alterations that may precede overt cardiomyocyte damage, offering potential avenues for biomarker development and therapeutic discovery. Top-down analysis of myosin light chains captures cell-specific heterogeneity and PTMs that may reflect contractile dysfunction and pathological remodeling. This molecular resolution provides mechanistic insight and reveals potential therapeutic targets. The role of succinylation in the heart, particularly in the context of heart failure and myofibrillar mechanics, remains poorly understood. This PTM reflects metabolic and mitochondrial dysregulation, key drivers of heart failure. It, therefore, represents a potential biomarker.^40,45^ Overall, proteoforms have been shown to play a critical role in numerous human diseases, including heart failure.^40^ Our SC-TDP platform provides a transformative tool for directly measuring intact proteins and their proteoforms in SCs. It enables detailed investigation of cellular function, intercellular variability, and disease mechanisms in the heart.

## METHODS

### Animal studies

All animal studies were approved by the Institutional Animal Care and Use Committee at Cedars-Sinai Medical Center and The Scripps Research Institute, and carried out in accordance with the National Institute of Health guidelines. Wild-type male and female mice (strain C57BL/6J) were purchased from Jackson Laboratories and were housed in colony cages (maximum of 2 animals per cage) with 12-hour shifts of the light-dark cycle. Animals were fed a standard chow diet (2018 Teklad Global 18% Protein diet, non-autoclavable, Inotiv).

### Protein standard Mixture

The Pierce Intact Protein Standard Mix (76 μg, Thermo FisherScientific, A33526) was reconstituted in Milli-Q water. Ten-microliter aliquots were prepared and stored at −80°C until use. For LC–MS analysis, an aliquot was thawed and diluted to a final concentration of 10 ng/μL in 30% trifluoroethanol (TFE) and 65% dimethyl sulfoxide (DMSO) in 5% formic acid.

### Extraction and Analysis of Bulk Cardiomyocyte Sample

Left ventricles were harvested from an adult male C57BL/6J wild-type mouse and processed for protein extraction. Fifty milligrams of left ventricular tissue were resuspended in 100 μL of ice-cold lysis buffer containing 30% TFE and 65% DMSO in 5% formic acid, supplemented with protease and phosphatase inhibitor cocktails (Thermo Fisher Scientific). Proteins were extracted from the heart tissue by mechanical disruption using a Dounce homogenizer until the tissue was fully homogenized. Cellular debris were pelleted by centrifugation (20,000g for 15□min at 4□°C) and the supernatant, which contained proteins, was transferred to a clean microcentrifuge tube and quantified using a BCA assay kit (Thermo Fisher Scientific). The lysate was immediately analyzed by LC-MS.

### Preparation and Analysis of Single Cardiomyocytes

Adult mouse ventricular cardiomyocytes were isolated as described previously.^46^ Male and female mice (8-12-week-old) were anesthetized using Nembutal via intraperitoneal injections (100 mg/kg). Hearts were excised and perfused retrogradely using the Langendorff preparation. Hearts were perfused for 15-18 min at 37°C with HEPES buffer containing 100 mg/mL collagenase type 2 (Cat#9001-12-1, Worthington, USA) and butanedione monoxime. At the end of the perfusion protocol, hearts were mechanically sheared and filtered through a 100-mm mesh filter (Cat#431752, Corning, Durham, NC). The single-cell suspension was centrifuged at 20 g for 3 minutes. For single-cell sorting, 200nL of purified water was dispensed per well in a 384-well plate. Plates were centrifuged (1500 rcf, 1 min, 4°C) and chilled at 4°C. Afterwards, cells were dispensed into wells and plates were centrifuged (1500rcf, 1min, 4°C). The 384-well plates were shipped on dry-ice to The Scripps Research Institute and stored at -80°C until use. The 384-well plate containing single cardiomyocytes in water were centrifuged for 5 min at 1,500 g (4°C). Each well was filled with 20µL of lysis buffer (30% TFE and 65% DMSO in 5% formic acid with protease and phosphatase inhibitor cocktails [Thermo Fisher Scientific]). The 384-well plate was centrifuged for 1 min at 1,500 g (4°C), sonicated in ice for ∼1 min, and then centrifuged for 1 min at 1,500 g (4°C).

### Nano LC Conditions

The standard protein mixture, bulk sample, and single cell lysates were analyzed using a nanocapillary Easy-nLC 1200 system coupled to an Orbitrap Fusion Lumos mass spectrometer (Thermo Fisher). The standard mixture, bulk sample, and single cardiomyocytes were separated on a fused silica capillary (∼20-30 cm × 75 μm i.d., with a 5 μm pulled tip) packed in-house with C4 reversed-phase resin (2.7 μm, 1000 Å, Halo). Mobile phase A was 95% H_2_O, 5% ACN in 0.1% formic acid and mobile phase B was 95% ACN, 5% H_2_O in 0.1% formic acid. Protein standards were eluted using a linear gradient from 25% to 55% solvent B in 35 min at a constant flow rate of 300 nL/min. The bulk sample and single cardiomyocytes were eluted using a linear gradient (60 minutes) which increased from 5 to 65% solvent B in the first 60 min followed by an increase to 95% solvent B over the next 1 min, and sustained 95% solvent B for the next 9 min at a constant flowrate of 300nL/min. The auto sampler temperature was set to 4°C. Both the standard protein mixture and bulk sample (∼10 ng [total protein]) were injected in triplicates.

### MS Conditions

The Orbitrap Fusion Lumos was operated in intact protein mode, with a nitrogen pressure of 2 mTorr in the ion routing multipole. The in-source fragmentation voltage was set to 15V. The precursor and fragment ions were acquired with a resolving power between 15,000 and 60,000. The automatic gain control (AGC) target values for MS and MS/MS scans were defined as 1E6 ions in 100 and 1000 ms, respectively. MS and MS/MS spectra were acquired by averaging 25 and 5 µscans, respectively. The RF lens was set to 75%. MS/MS spectra were generated in data-dependent acquisition (top N) where the two most abundant precursors were isolated by the quadrupole with a 3 m/z isolation window. Precursor ions were dynamically excluded for 6000 seconds after being selected. Two fragmentation modes were used in consecutive scans: 1) electron-transfer/higher-energy collision dissociation (EThcD) with calibrated ETD charge-dependent parameters and supplemental activation (HCD collision energy of 23-25%). The MS data from bulk sample were acquired only using EThcD.

### Data Analysis

Raw data from the standard protein mixture were processed using Freestyle (Thermo Fisher Scientific) and UniDec.^47^ Raw files (bulk sample and individual cardiomyocytes) were processed using ProSight PD 4.5 (Thermo Fisher Scientific). Precursor and fragment ions were deconvoluted using the Xtract algorithm and then searched against a Mus Musculus database (Uniprot) containing annotated modifications. The database was subsequently converted into a ProSight PD-compatible (.psdb) database. All searches were performed using a three-node strategy, with data acquired from the EThcD and HCD dissociation modes searched simultaneously. Proteins and proteoforms were identified as follows: 1) complete sequences with unexpected modifications were identified using a wide absolute precursor mass tolerance (200 Da); 2) known (well-matching) proteoforms with complete sequences were identified using a narrow absolute precursor mass tolerance (2.2 Da); and 3) truncated proteoforms were identified using a narrow precursor biomarker mass tolerance (10 ppm). For all three nodes, the fragment mass tolerance was set to 10 ppm. Results were filtered using a false discovery rate (FDR) threshold of 1% at both the protein and proteoform levels.^48^ ProSight Lite and TDValidator in ProSight PD 4.5 software were used to provide graphical interpretation of tandem MS spectra, in which inter-residue cleavages were indicated in blue for *b* and *y* ions and in red for *c* and *z* ions.

## ASSOCIATED CONTENT

Support information.

## AUTHOR INFORMATION

### Author Contributions

The manuscript was written through contributions of all authors. All authors have given approval to the final version of the manuscript.

### Notes

K.R.D. is involved in the commercialization of the software ProSight PD 4.5. The other authors declare no conflict of interest.

### Data availability

Raw spectrum data are available via figshare.

## ACKNOWLEDGMENTS

This work was supported by the ORAU Ralph E. Powe Junior Faculty Enhancement Award (F.P.G.), the National Institutes of Health (NIH), the National Heart, Lung and Blood Institute (R00-HL141702, R01-HL177461 to A.K.), the Leukemia Research Foundation New Investigator Award (grant No. 941997 to A.K.), the 1R01HL155346-01A1 (J.E. Van Eyk), 1R01HL144509-01 (J.E. Van Eyk), The Erika Glazer Endowed Chair (J.E. Van Eyk), 1R21MH129776-01 (J.R.Y), R01 MH100175-05 (J.R.Y), and R01 DK138430 (E.S.).

