## Supplemental Figures for "Capturing Cardiomyocyte Cell-to-Cell Heterogeneity via Shotgun Single Cell Top-Down Proteomics"

### Fabio P. Gomes*^[a]^, Blandine Chazarin^[b,c]^ , Aleksandra Binek^[b,c]^, Aleix Navarro Garrido^[d]^, Kenneth R. Durbin^[e]^, Ricard Garcia-Carbonell^[d]^, Kanchan Pathak^[a]^, Delaynie Brinkman^[a]^, Reynaldo Magalhaes Melo^[d]^, Enrique Saez^[d]^, Anja Karlstaedt^[c]^[, Jennifer E. Van Eyk](https://pubmed.ncbi.nlm.nih.gov/?term=Van+Eyk+JE&cauthor_id=33067450)^[b,c]^, and John R. Yates III*^[d]^

Virginia Commonwealth University, Department of Chemistry, Richmond, VA 23284 ^[a]^; Advanced Clinical Biosystems Research Institute, The Smidt Heart Institute, Cedars-Sinai Medical Center, Los Angeles, CA 90048 ^[b]^; Department of Cardiology, The Smidt Heart Institute, Cedars-Sinai Medical Center, Los Angeles, CA 90048 ^[c]^, The Scripps Research Institute, Departments of Integrative Structural and Computational Biology and Molecular and Cellular Biology, La Jolla, CA 92037 ^[d]^; Proteinaceous, Evanston, IL 60201 ^[e]^

Corresponding Authors

*(F.P.G)

*(J.R.Y III)

**Table of Contents**

**Supplementary Figures**

- **Figure S1.** Total ion chromatograms (TICs) of intact protein standard mixtures (~9 - 68 kDa) prepared by diluting the samples in lysis buffer (30% TFE and 65% DMSO in 5% formic acid with protease and phosphatase inhibitor cocktails, three overlapped replicates).
- **Figure S2.** TICs of a bulk cardiomyocyte sample (10 ng total protein) isolated from mouse heart tissue and lysed using lysis buffer (30% TFE and 65% DMSO in 5% formic acid with protease and phosphatase inhibitor cocktails, triplicate measurements). The time ranges in which each of the two selected proteins or proteoforms were detected are highlighted using different colors.
- **Figure S3.** Mass distribution of identified proteoforms in the bulk sample (~2 - 22 kDa) across all replicates.
- **Figure S4. Overlap of proteins and proteoforms across three technical replicates from a bulk cardiomyocyte sample.** Venn diagrams show the number of shared and unique identifications. Proteins were matched based on accession numbers. Proteoforms were compared based solely on protein descriptions and associated modifications, so the identification numbers may not represent the total number of proteoforms listed in **Table S1**.
- **Figure S5. Schematic of the TDP workflow for the identification and characterization of proteoforms from single cardiomyocytes. A)** The heart is extracted and dissected from an adult male wild-type mouse. **B)**  Cardiomyocytes are isolated from the mouse heart. **C)** The cardiomyocyte suspension is transferred to the CellenOne X1 device, which dispenses individual cardiomyocytes into a 384-well plate. **D)** Proteins are extracted from individual cardiomyocytes directly within the wells of the 384-well plate. **E)** Protein extracts from individual cardiomyocytes are analyzed by LC–MS/MS using both EThcD and HCD fragmentation methods. **F)** The MS1 scan detects intact proteoforms. **G)** The MS2 scan fragments MS1-selected proteoforms using either EThcD or HCD for characterization. **H)** Proteoform identification is performed using ProSight PD 4.5 software with a false discovery rate (FDR) cutoff of 1%. Figures were created with BioRender.com.
- **Figure S6.** Percentage of proteoforms identified using EThcD and HCD across 13 individual cardiomyocytes with a comparative contribution of each fragmentation method.
- **Figure S7.** **Overlap of proteins and proteoforms identified in the 13 individual cardiomyocytes and across all three replicates of the bulk cardiomyocyte tissue sample**. Proteins were compared based on accession numbers. To facilitate comparisons, proteoforms were matched exclusively based on protein descriptions and associated modifications; therefore, the identification numbers may not represent the total number of proteoforms listed in **Tables S1** and **S2**.
- **Figure S8.** Mass distribution of identified proteoforms from 13 individual cardiomyocytes (~2 - 22 kDa).
- **Figure S9**. Fragmentation map of the trimethylated and phosphorylated MLC-2 proteoform lacking phosphorylation at residue T51, generated using ProSight Lite.
- **Figure S10. Fragmentation maps of MYL3 proteoforms.** **A)** Trimethylated MYL3 proteoform. **B)** MYL3 proteoform without containing combinatorial novel trimethylation at A1 and known phosphorylation at residue S187.
- **Figure S11**. Fragmentation map of the COX6c proteoform with one acetylation at the N-terminus.
- **Figure S12.** Fragmentation map of the Qcr7 proteoform bearing a succinylation at K11, with diagnostic ions *y109 (+16), y103 (+14),* and *b90 (+12)*.

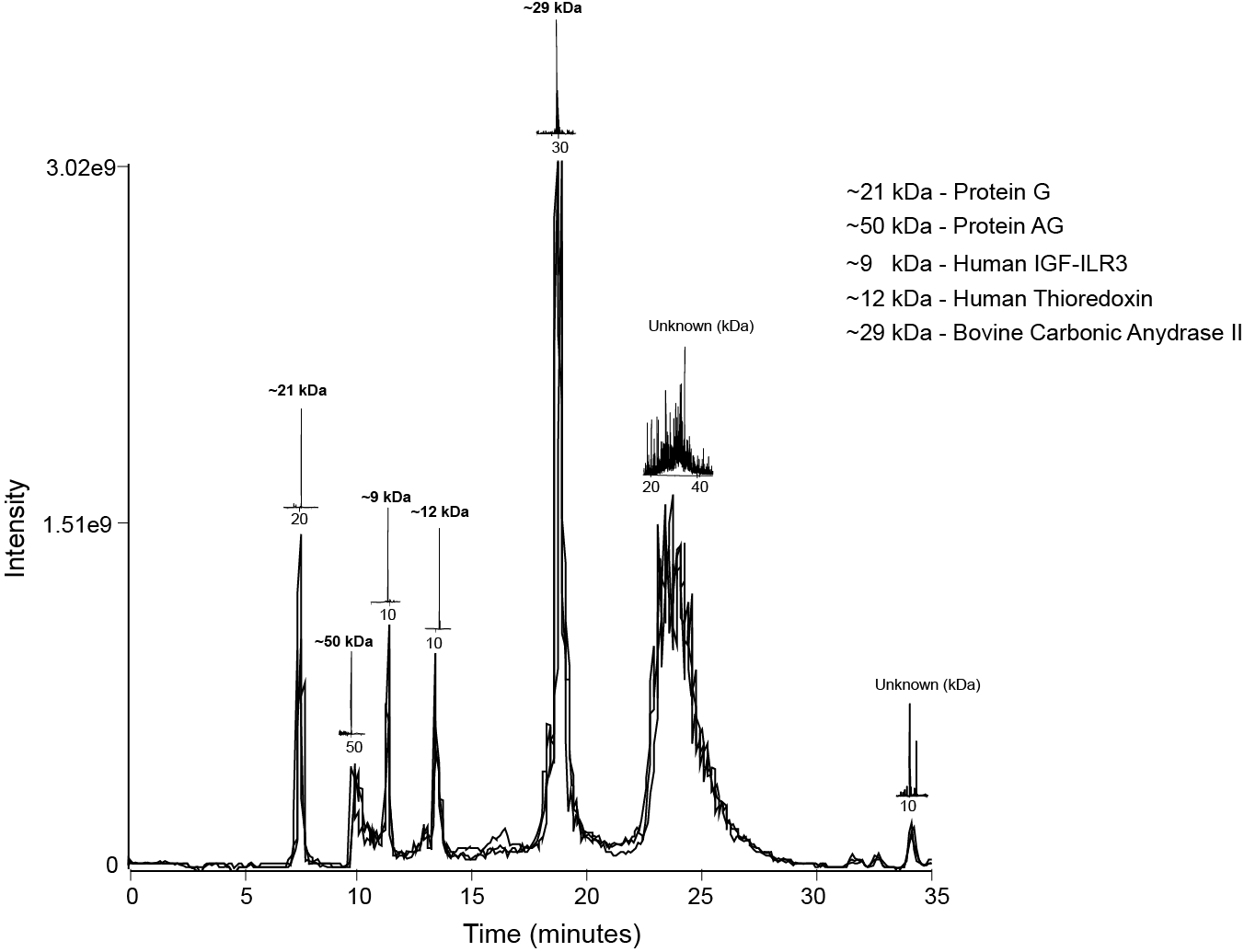

**Figure S1.** Total ion chromatograms (TICs) of intact protein standard mixtures (~9 - 68 kDa) prepared by diluting the samples in lysis buffer (30% TFE and 65% DMSO in 5% formic acid with protease and phosphatase inhibitor cocktails, three overlapped replicates).

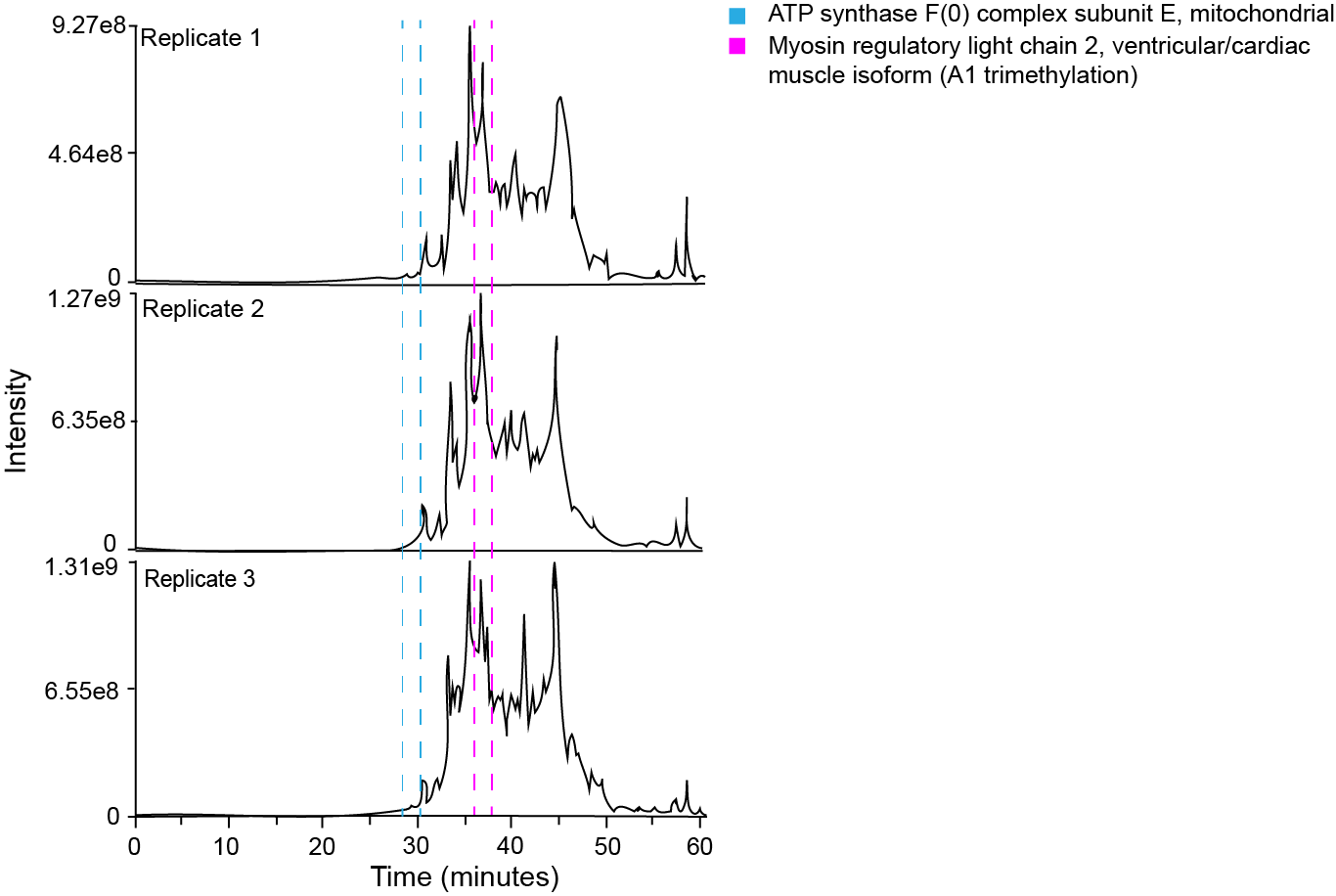

**Figure S2.** TICs of a bulk cardiomyocyte sample (10 ng total protein) isolated from mouse heart tissue and lysed using lysis buffer (30% TFE and 65% DMSO in 5% formic acid with protease and phosphatase inhibitor cocktails, triplicate measurements). The time ranges in which each of the two selected proteins or proteoforms were detected are highlighted using different colors.

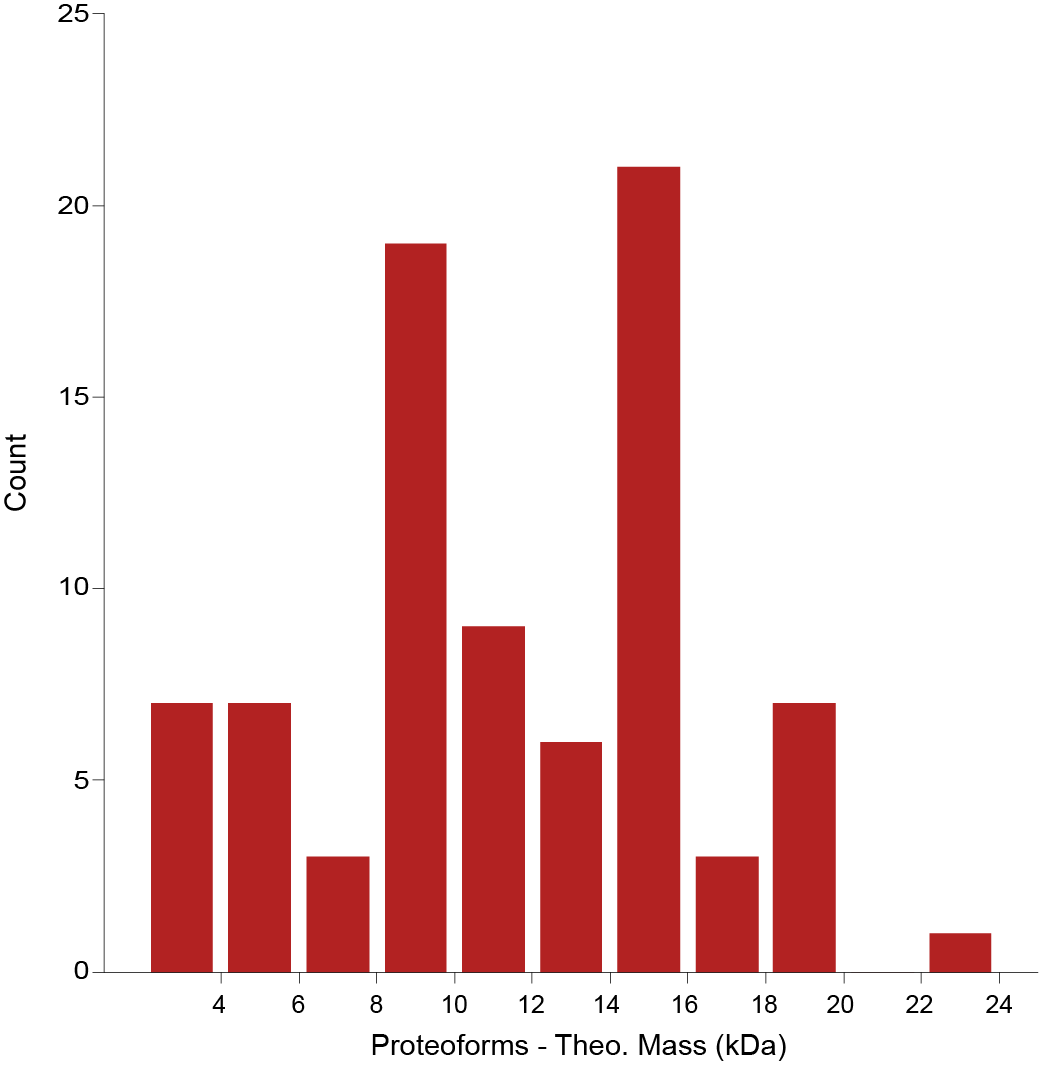

**Figure S3.** Mass distribution of identified proteoforms in the bulk sample (~2 - 22 kDa) across all replicates.

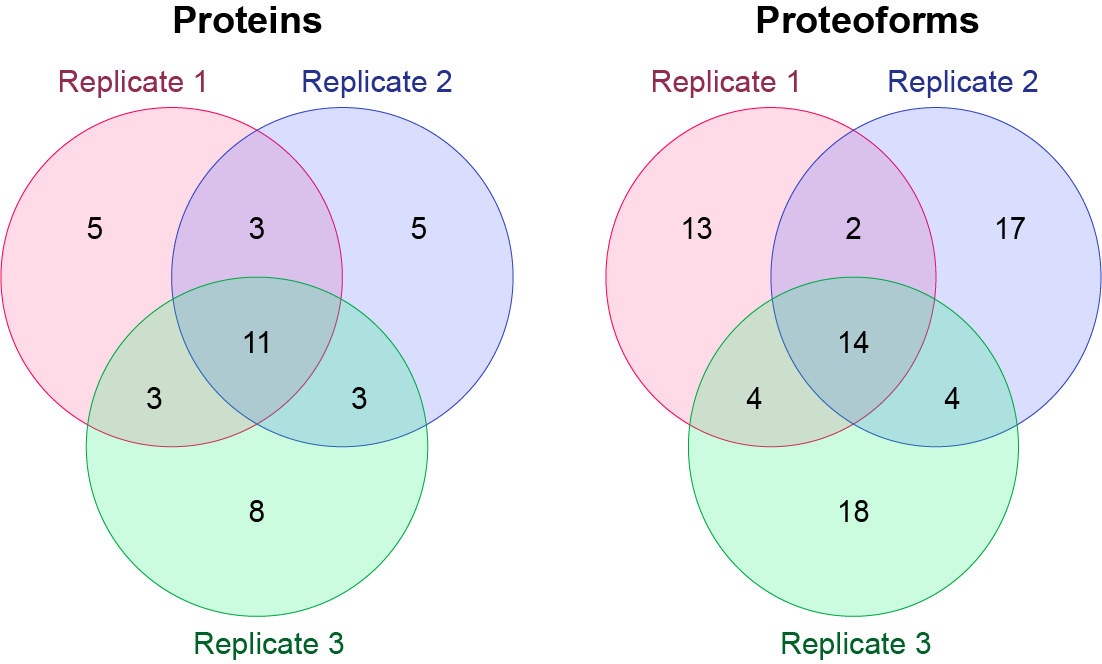

**Figure S4. Overlap of proteins and proteoforms across three technical replicates from a bulk cardiomyocyte sample.** Venn diagrams show the number of shared and unique identifications. Proteins were matched based on accession numbers. Proteoforms were compared based solely on protein descriptions and associated modifications, so the identification numbers may not represent the total number of proteoforms listed in **Table S1**.

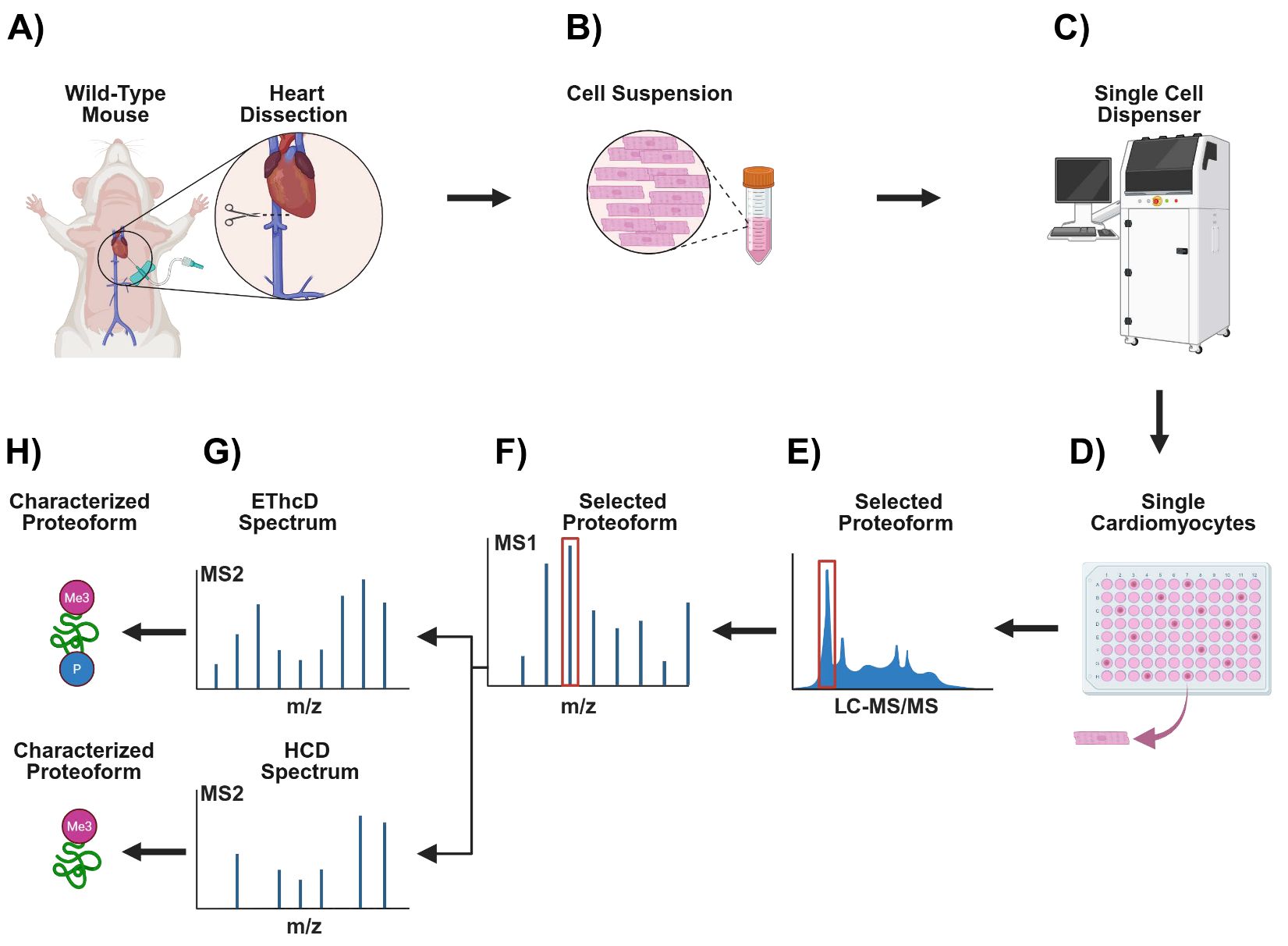

**Figure S5. Schematic of the TDP workflow for the identification and characterization of proteoforms from single cardiomyocytes. A)** The heart is extracted and dissected from an adult male wild-type mouse. **B)**  Cardiomyocytes are isolated from the mouse heart. **C)** The cardiomyocyte suspension is transferred to the CellenOne X1 device, which dispenses individual cardiomyocytes into a 384-well plate. **D)** Proteins are extracted from individual cardiomyocytes directly within the wells of the 384-well plate. **E)** Protein extracts from individual cardiomyocytes are analyzed by LC–MS/MS using both EThcD and HCD fragmentation methods. **F)** The MS1 scan detects intact proteoforms. **G)** The MS2 scan fragments MS1-selected proteoforms using either EThcD or HCD for characterization. **H)** Proteoform identification is performed using ProSight PD 4.5 software with a false discovery rate (FDR) cutoff of 1%. Figures were created with BioRender.com.

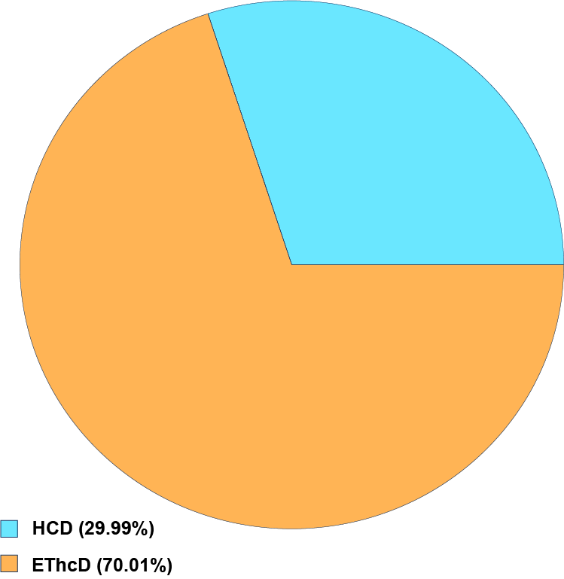

**Figure S6.** Percentage of proteoforms identified using EThcD and HCD across 13 individual cardiomyocytes with a comparative contribution of each fragmentation method.

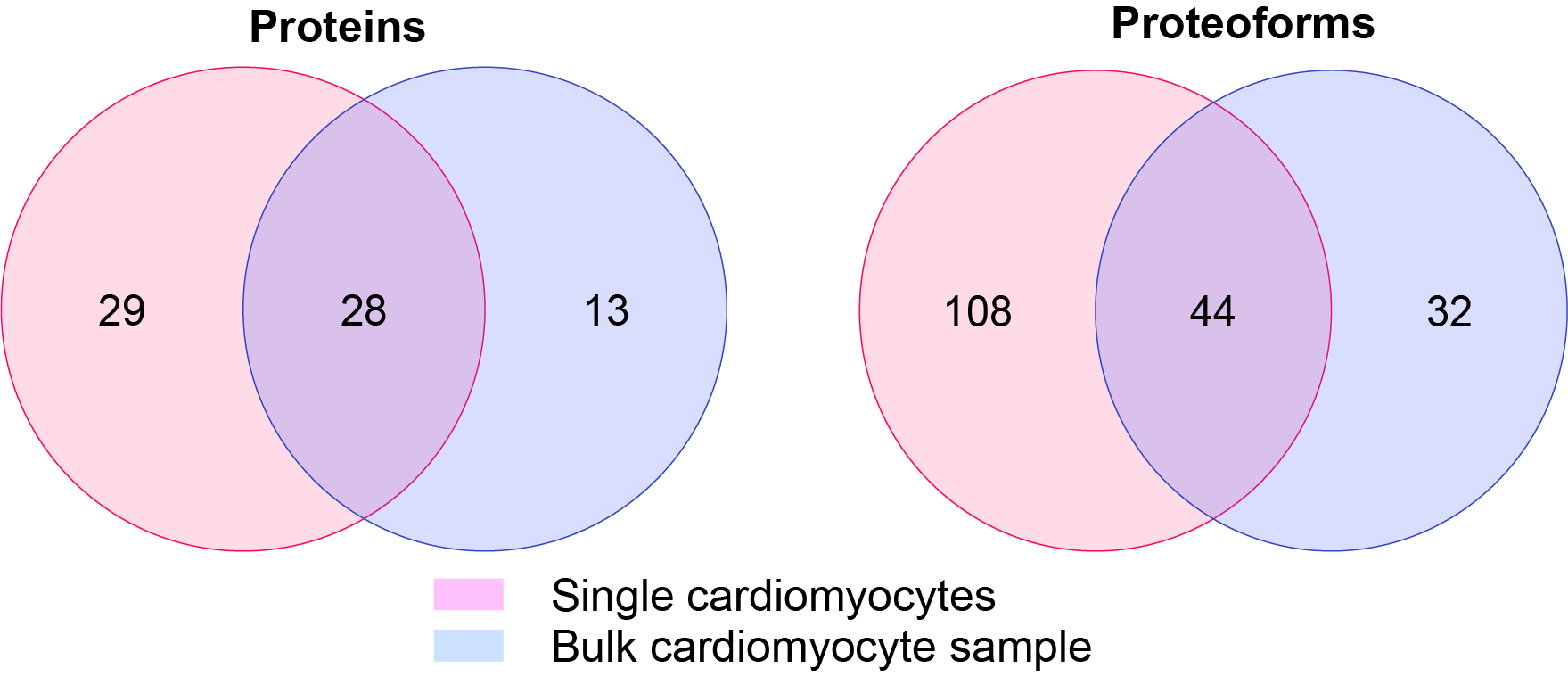

**Figure S7.** **Overlap of proteins and proteoforms identified in the 13 individual cardiomyocytes and across all three replicates of the bulk cardiomyocyte tissue sample**. Proteins were compared based on accession numbers. To facilitate comparisons, proteoforms were matched exclusively based on protein descriptions and associated modifications; therefore, the identification numbers may not represent the total number of proteoforms listed in **Tables S1** and **S2**.

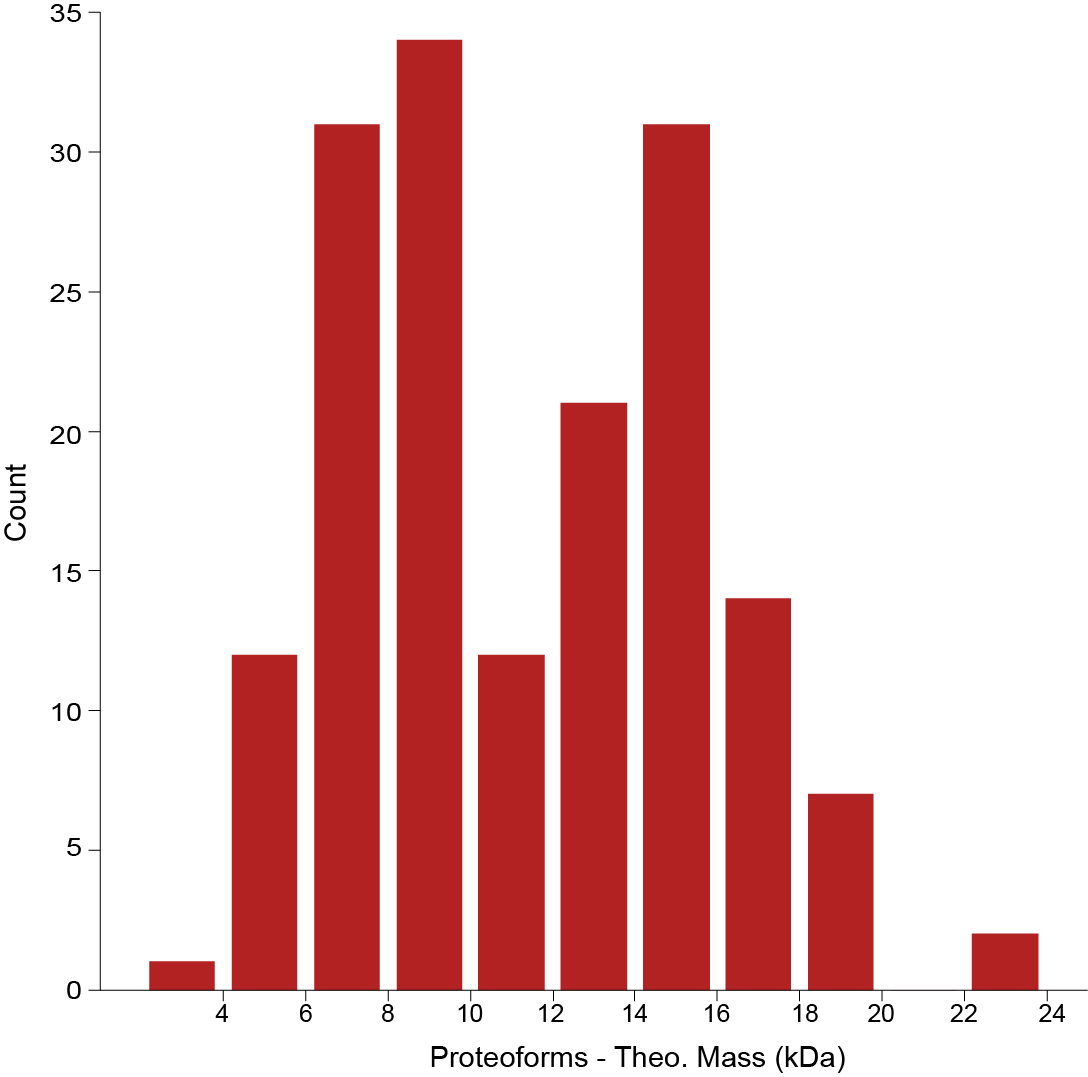

**Figure S8.** Mass distribution of identified proteoforms from 13 individual cardiomyocytes (~2 - 22 kDa).

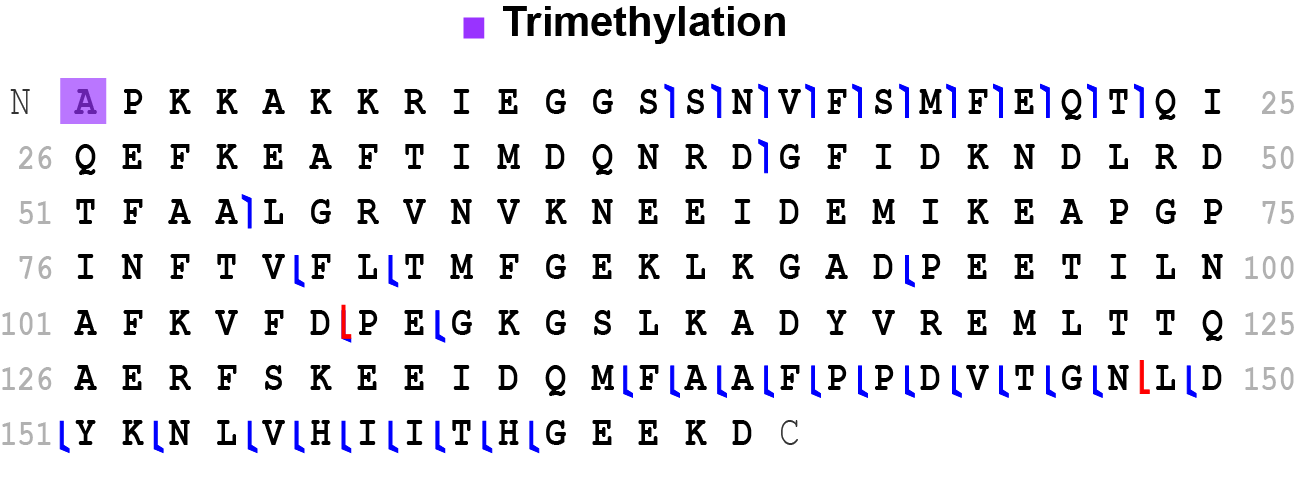

**Figure S9**. Fragmentation map of the trimethylated and phosphorylated MLC-2 proteoform lacking phosphorylation at residue T51, generated using ProSight Lite.

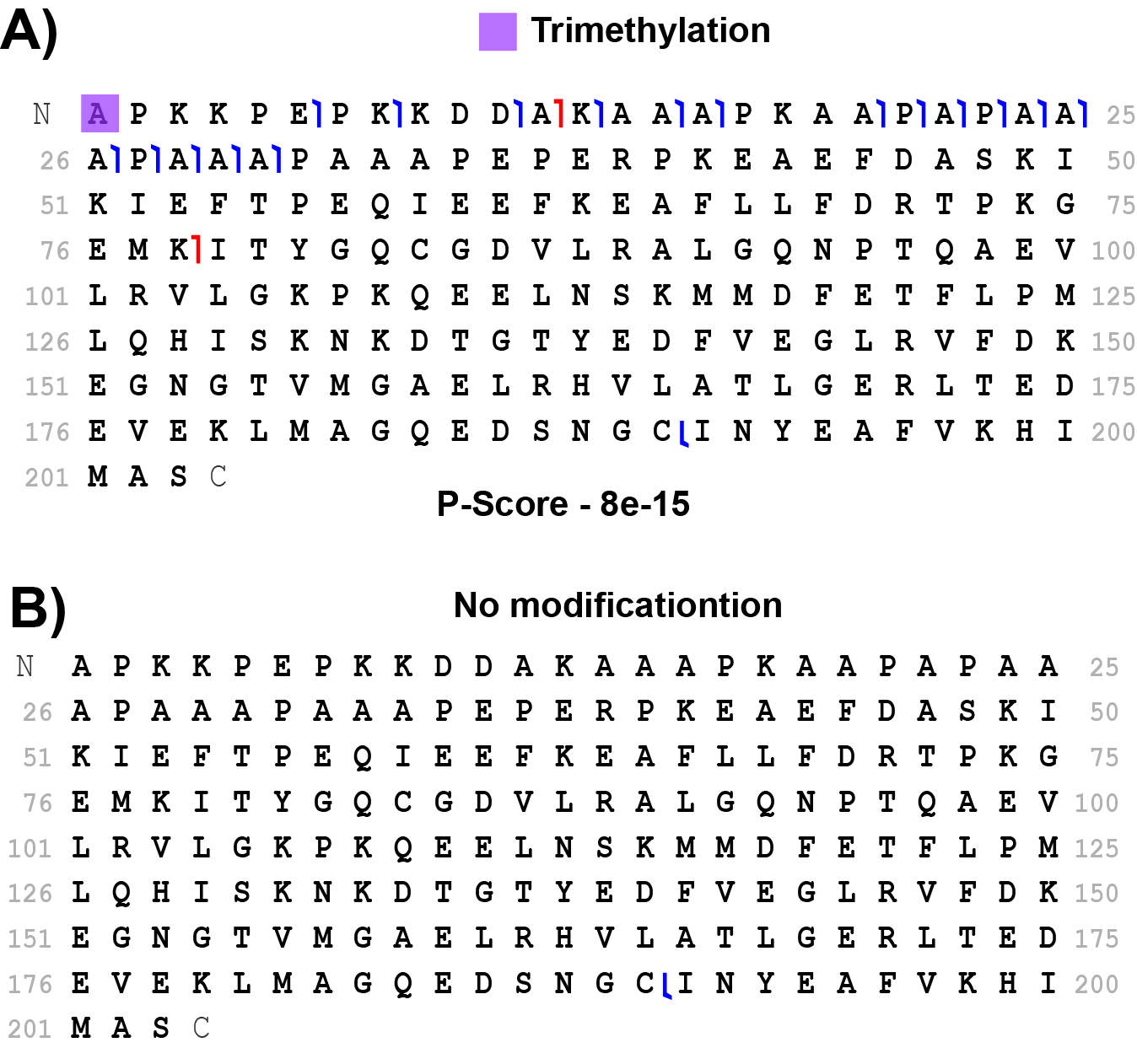

**Figure S10. Fragmentation maps of MYL3 proteoforms.** **A)** Trimethylated MYL3 proteoform. **B)** MYL3 proteoform without containing combinatorial novel trimethylation at A1 and known phosphorylation at residue S187.

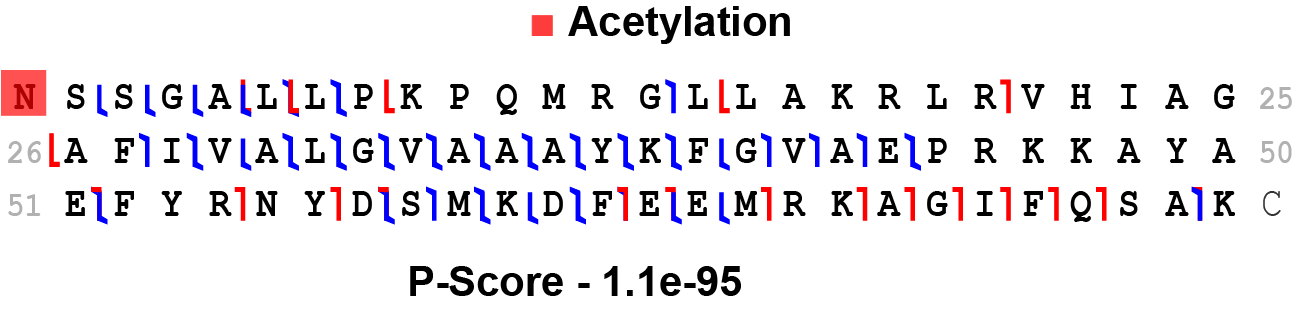

**Figure S11**. Fragmentation map of the COX6c proteoform with one acetylation at the N-terminus.

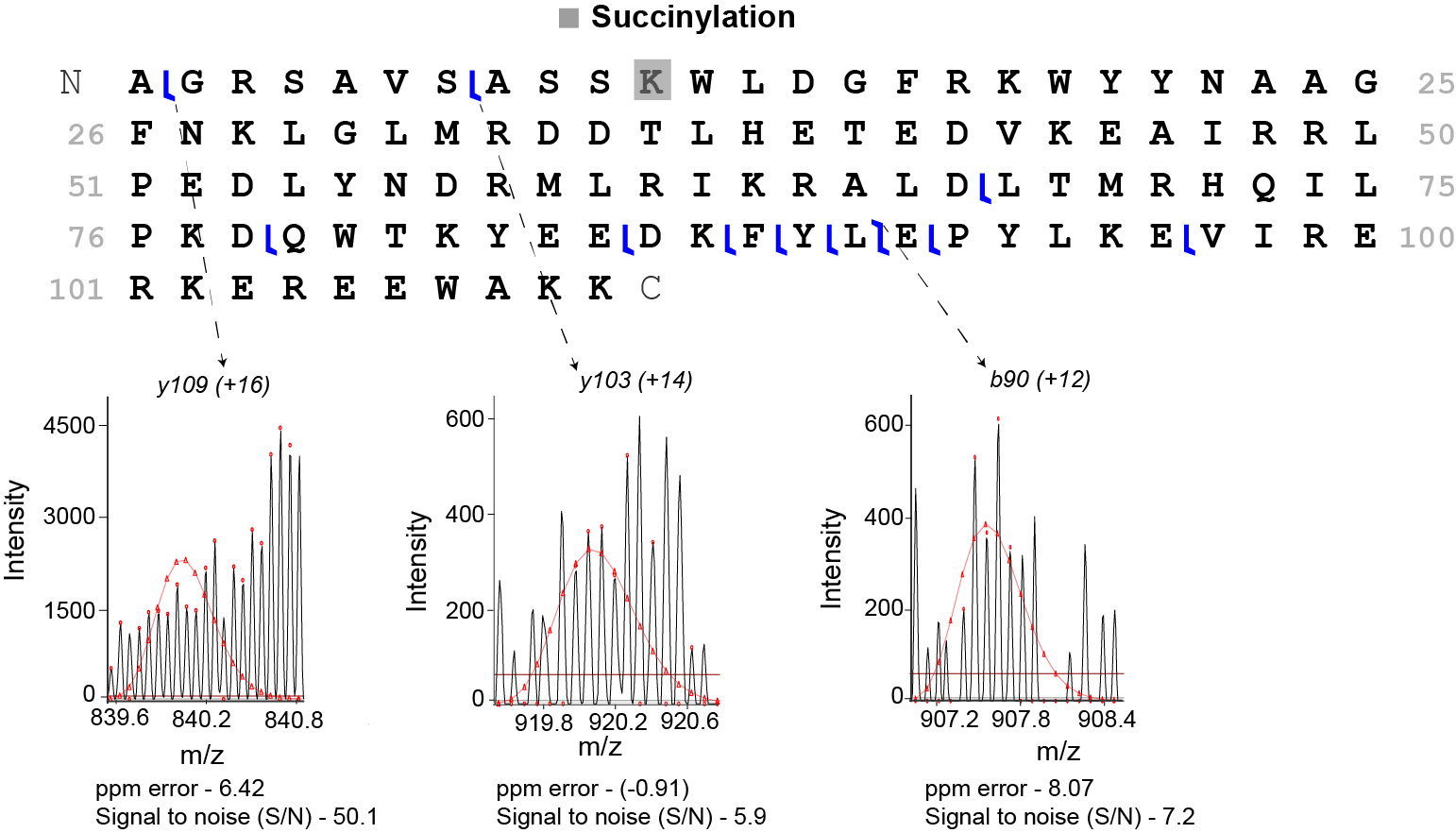

**Figure S12.** Fragmentation map of the Qcr7 proteoform bearing a succinylation at K11, with diagnostic ions *y109 (+16), y103 (+14),* and *b90 (+12)*.
